# Cell-nonautonomous local and systemic responses to cell arrest enable long-bone catch-up growth in developing mice

**DOI:** 10.1101/218487

**Authors:** Alberto Roselló-Díez, Linda Madisen, Sébastien Bastide, Hongkui Zeng, Alexandra L. Joyner

## Abstract

Catch-up growth after insults to growing organs is paramount to achieving robust body proportions. In fly larvae, local injury is followed by local and systemic compensatory mechanisms that allow damaged tissues to regain proportions with other tissues. In vertebrates, local catch-up growth has been described after transient reduction of bone growth, but the underlying cellular responses are controversial. We developed an approach to study catch-up growth in foetal mice by inducing mosaic expression of the cell cycle suppressor p21 in the cartilage cells (chondrocytes) that drive long bone elongation. By specifically targeting the left hindlimb, the right limb served as an internal control. Strikingly, left-right limb symmetry was not altered, revealing deployment of compensatory mechanisms. Above a certain threshold of insult, an orchestrated response was triggered involving local enhancement of bone growth and systemic growth reduction that ensured body proportions were maintained. The local response entailed hyper-proliferation of spared left-limb chondrocytes that was associated with reduced chondrocyte density. The systemic effect involved impaired placental IGF signalling and function, revealing bone-placenta communication. Thus, vertebrates, much like invertebrates, can mount coordinated local and systemic responses to developmental insults to ensure normal body proportions are maintained.

## Introduction

An important question in biology is how cells integrate intrinsic and extrinsic information such that their combined behaviours produce higher-order processes and structures, as seen during organogenesis and tissue repair. In *Drosophila* larvae, injured tissues can undergo compensatory proliferation[1] as well as secrete an alarm signal that triggers both a systemic developmental delay and growth reduction[2–5]. Together, these processes allow the damaged tissue(s) to catch-up with other tissues, but the role of damaged vs. undamaged cells remains controversial[6,7]. In vertebrates, systemic growth reduction after injury in a non-essential organ has not been reported. However, systemic catch-up growth has been described after transient impairment of whole-body growth[8–10], and local growth compensation can occur after unilateral manipulation of long bones within the limbs[11]. Tight control of inter-limb and limb-body proportions are critical for efficient locomotion and interaction with the environment, and therefore long bones are an excellent model for studies of growth regulation. Growth of the initial cartilage templates of long bones is driven by the growth plates (GPs) at each end, where chondrocytes proliferate, then mature, become hypertrophic and eventually are replaced by bone-forming cells in a process called endochondral ossification[12]. It has been proposed that bone catch-up growth is due to a cell-autonomous delay in the normal developmental decline of chondrocyte proliferation, such that when the insult is lifted, the formerly arrested chondrocytes retain a higher proliferative potential[9,13]. A similar mechanism was suggested to apply to other organs as well[14]. However, such a mechanism does not account for cases in which catch-up growth is faster than expected for the observed maturation delay[15,16]. Here, we developed new mouse models to transiently decrease long-bone growth in mice in order to determine the contributions of cell-autonomous and nonautonomous regulation during catch-up growth.

## Results and Discussion

### Mosaic local proliferation blockade in the left limb cartilage does not lead to a major left-specific bone growth reduction

A major roadblock for studies of intra- and inter-organ growth regulation in mouse embryos has been a lack of models in which growth rate can be altered in a specific cell type within an organ, and ideally in only one of two paired organs, leaving the unmanipulated organ as an internal control. To address this deficiency, we devised new mouse models of inducible and transient growth inhibition in the left limb. We generated an *Igs7^TRE-LtSL-p21/+^* allele, a variant of a double-conditional allele[17], to achieve doxycycline (Dox)-tuneable misexpression of the cell cycle suppressor *Cdkn1a* (*p21* hereafter)[18] in the cells where Cre and (r)tTA activity intersect (Fig. 1A-B). Due to a floxed tdTomato-STOP sequence (LtSL), expression of tdTomato (tdT) takes place in cells expressing (r)tTA but having no history of Cre activity, whereas *p21* is expressed in the cell population with a history of Cre and current (r)tTA activity (Fig. 1A). We named the general type of allele Doxycycline-controlled and Recombinase Activated Gene OverexpressioN (DRAGON). By combining the *DRAGON-p21* allele with an *asymmetric-Pitx2-enhancer-Cre* line expressing Cre in the precursors of the left limb mesenchyme[19] (Supplemental Fig. 1A-F) and a *Col2a1-rtTA* line[20] (Fig. 1B), Dox-dependent ectopic *p21* expression was achieved specifically in non-hypertrophic chondrocytes of the left limb cartilage elements (Fig. 1C-C’). Consequently, any growth adjustment detected in the right limb of triple transgenic animals (*Pit-Col-p21*) when compared to control littermates must be due to activation of a systemic effect.

**Figure 1.**
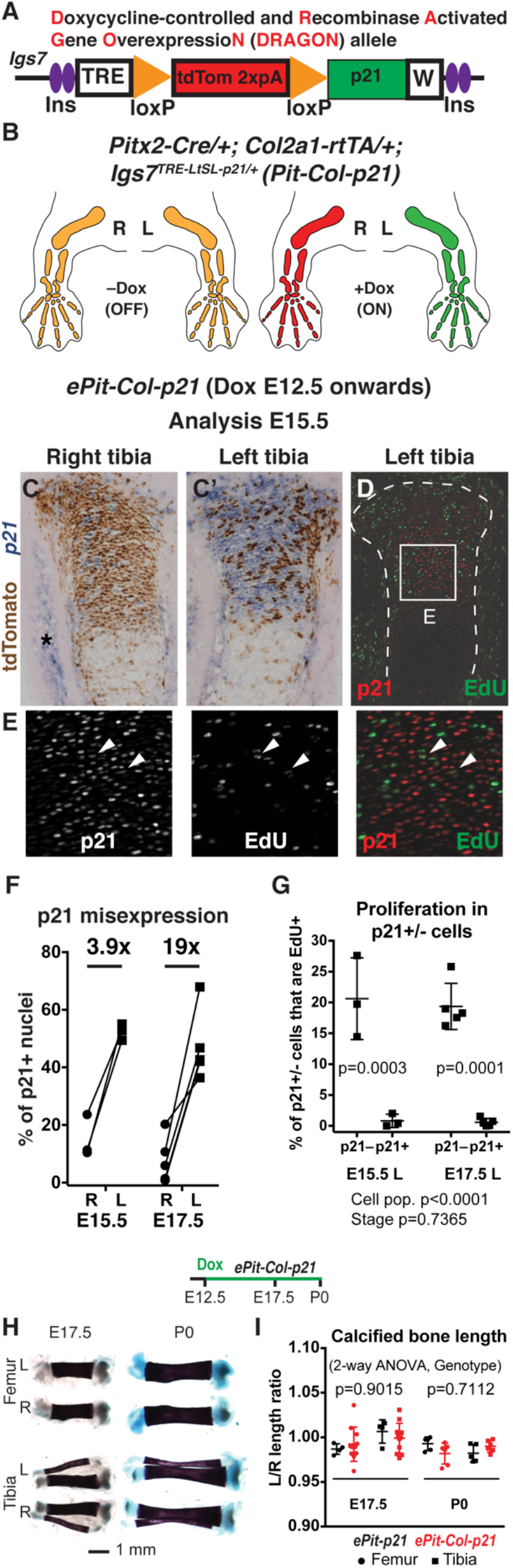
Mosaic local proliferation blockade in the left limb cartilage does not lead to a major left-specific bone growth reduction. (**A**) *DRAGON-p21* allele in the *Igs7* locus. Ins=insulator, TRE=Tetracycline-responsive element, 2xpA=transcriptional STOP, W=WPRE (mRNA-stabilizing sequence) followed by pA. (**B**) Schematic showing *p21* expression driven by the left-specific *Pitx2-Cre* and cartilage-specific *Col2a1-rtTA* (*Pit-Col-p21*). (**C-E**) Expression of tdT protein and *p21* mRNA (C, C’), and p21 protein and EdU (D, E) at E15.5, with Dox administered at E12.5. n=3. Box in (D) is magnified in (E). Asterisk=endogenous *p21* expression. Arrowheads=rare double-positive cells. (**F**) Quantification of p21^+^ cells in the proliferative zone of *ePit-Col-p21* proximal tibias, at E15.5 (n=3) and E17.5 (n=5). The average left/right fold-change is indicated. (**G**) Quantification of EdU incorporation in p21^+^ and p21^−^ cells of left *ePit-Col-p21* proliferative zone of the cartilage, at E15.5 and E17.5 (n=3 and 5). Comparison by 2-way ANOVA with Cell population and Stage as variables (p-values below graphs). p-values for Sidak’s multiple comparisons posthoc test (between Cell populations) are shown on the graph. (**H, I**) Skeletal preparations (H) and quantification of the left/right ratio [(I), mean±SD] of the calcified region of femur and tibia at E17.5 (n=4 *ePit-p21* and 11 *ePit-Col-p21* mice) and P0 (n=5 and 6). At each stage, data were analysed by 2-way ANOVA with Genotype and Bone identity as variables. p-values for Genotype are shown.

When Dox was administered from embryonic day (E) 12.5 until birth (*ePit-Col-p21* model), analysis at E14.5-E17.5 revealed the expected cartilage-exclusive expression of tdT, mainly in the right skeletal elements, and *p21* expression preferentially in the left limb cartilage, albeit in a mosaic fashion (Supplemental Fig. 1G-H, Fig. 1C-F; 36-67% vs. 0.8-23% of chondrocytes were p21+ in left vs. right proximal tibia). Since Cre activity and therefore *p21* expression was more widespread in the left hindlimb than in the left forelimb (Supplemental Fig. 1I-J and [21]), we focused our initial analysis on the hindlimb. As expected, proliferation was inhibited in p21+ chondrocytes at E15.5 and E17.5 (Fig. 1D-E and G). Importantly, misexpression of *p21* in proliferative zone (PZ) chondrocytes did not induce precocious expression of chondrocyte maturation markers (e.g. *Ihh, Col10a1, Cdkn1c*) or cell senescence (monitored by expression of p16) by E17.5, nor did it alter the archetypical cytoarchitecture of the cartilage[12] or chondrocyte survival at E15.5 or E17.5 (Supplemental Fig. 2A-G). However, the normal expression domains of *Ihh, Cola10a1* and *Cdkn1c* in (pre)hypertrophic chondrocytes (which no longer expressed the transgene) appeared slightly fainter in the left cartilage, suggesting a mild maturation impairment (Supplemental Fig. 2C-E).

Strikingly, *ePit-Col-p21* mice at E17.5 or birth (P0) showed no obvious differences in the left/right ratio of tibia and femur length compared to *Pitx2-Cre; Igs7^TRE-LtSL-p21/+^* control littermates (*ePit-p21*) (Fig. 1H-I), indicating that compensatory mechanisms had been activated to maintain body proportions. Hereafter, we refer to this new type of catch-up growth that happens during an on-going insult as ‘adaptive growth’.

### Cell-nonautonomous compensation by spared neighbours in response to mosaic blockade of chondrocyte proliferation

To elucidate the compensatory mechanisms underlying adaptive growth, we first tested for an organ-intrinsic response, focusing on a change in chondrocyte proliferation. Indeed, the left/right ratio of EdU incorporation by p21^−^ chondrocytes was higher in experimental animals as compared with controls at E17.5 and P0 but not E15.5 (Fig. 2A and Supplemental Fig. 2H), revealing cell-nonautonomous compensatory proliferation of p21^−^ cells in the presence of p21^+^ neighbours. Since p21^−^ cells did not differ in size from those of control mice (Supplemental Fig. 2I), the hyperproliferation of these cells at E17.5 likely contributes to the lack of a left-specific growth reduction in *ePit-Col-p21* embryos. In fact, overall EdU incorporation in left and right *ePit-Col-p21* GPs (without distinguishing between p21^+^ and p21^−^ cells), while tending to be reduced was not significantly different, indicating that the compensatory proliferation phenomenon is quite effective (Fig. 2B). Moreover, the proliferative disadvantage of E17.5 p21^+^ vs. p21^−^ chondrocytes in the left limb of *ePit-Col-p21* mice resulted in dilution of p21+ chondrocytes from 45-50% of PZ chondrocytes at E15.5 and E17.5 to ~20% at P0 (Fig. 2C-D), and this depletion was not due to inactivation of rtTA activity (Fig. 2D). Our finding that a compensatory response occurs during the insult and involves cell-nonautonomous mechanisms is distinct from a model that proposes compensation is cell-autonomous once the insult is lifted[9,11,13], and thus introduces a new conceptual framework for the interpretation of previous and future results concerning long-bone growth.

**Figure 2.**
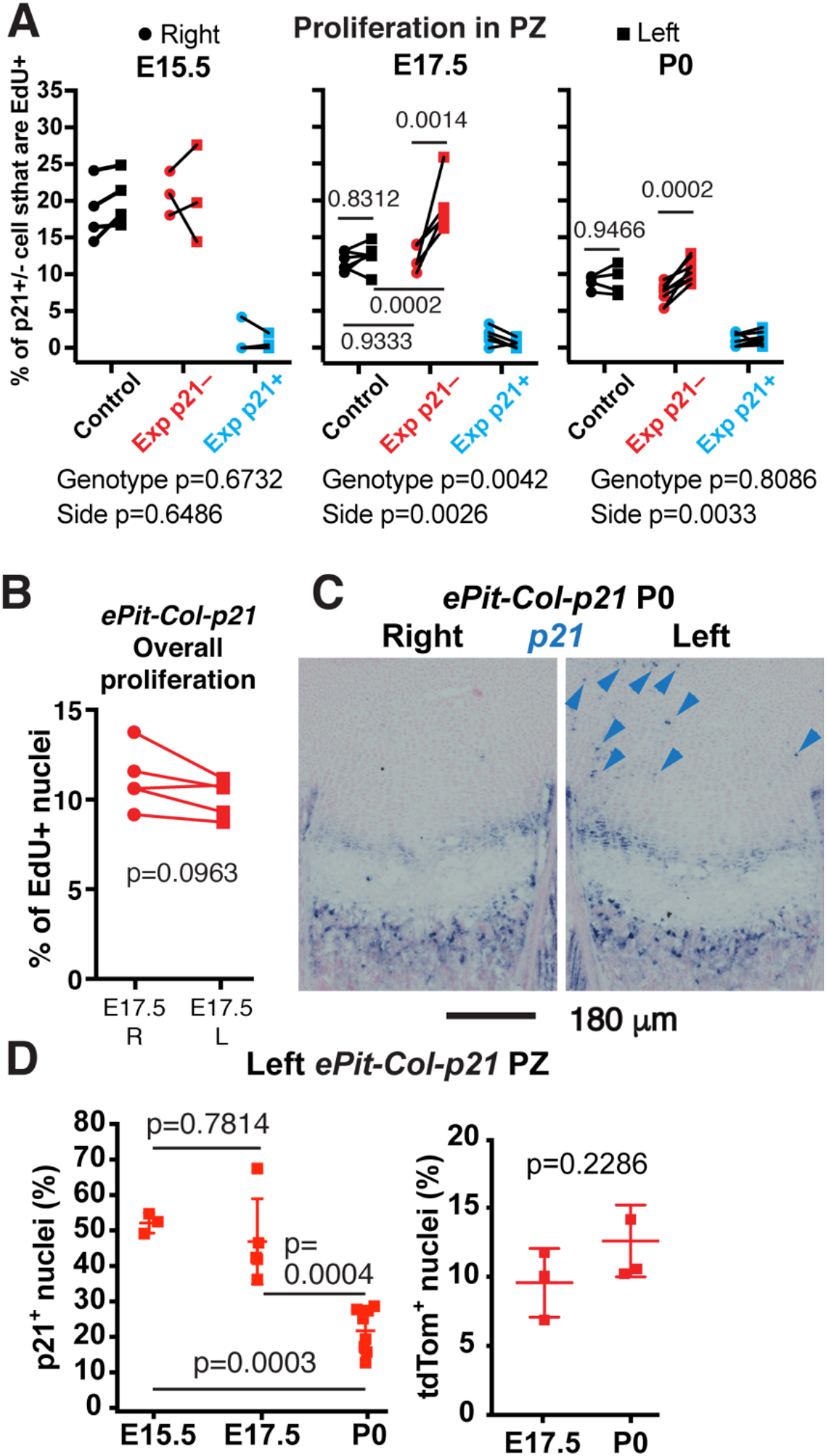
Cell-nonautonomous compensation by spared neighbours in response to mosaic blockade of chondrocyte proliferation. (**A**) % of p21^+^ or p21^−^ chondrocytes that have EdU^+^ nuclei in the proliferative zone (PZ) in the left and right tibias of E15.5, E17.5 and P0 *ePit-p21* (Control, n=4, 6 and 4) and *ePit-Col-p21* (Exp, n=3, 5 and 8) embryos. p21^−^ cells from Control and Exp mice were compared by 2-way ANOVA with Side and Genotype as variables (p-values below graphs). For each significant variable, p-values for Sidak’s multiple comparisons posthoc test are shown. (**B**) % of EdU^+^ chondrocytes in the PZ of left and right proximal tibias of E17.5 *ePit-Col-p21* embryos, without distinguishing by p21 expression. Comparison by paired two-tailed t-test. (**C-D**) *In situ* hybridisation of *p21* [(C), arrowheads denote ectopic expression] and quantification of tdT and p21 (D) on sections of left *ePit-Col-p21* tibial GPs at E15.5, E17.5 and P0. n=3, 5 and 8 for p21; 3 at each stage for tdT. The % of p21+ cells was compared by one-way ANOVA (p<0.0001). p-values for Tukey’s multiple comparisons posthoc test are shown. The % of tdT^+^ cells (a proxy for rtTA activity) was compared by unpaired two-tailed Mann-Whitney test.

### Compensatory proliferation takes place when cell density in the growth plate is lower than normal

To learn whether compensatory proliferation was independent of interactions with other tissues, we cultured left and right E15.5 *ePit-Col-p21* tibiae (together in the same well) for two days with Dox, in the absence of soft tissues (Fig. 3A). Notably, EdU incorporation in p21^−^ chondrocytes was significantly higher in the left as compared to the right cultured cartilage (Fig. 3B-C), indicating that compensatory proliferation is a cartilage-intrinsic phenomenon. We next addressed whether the amount of p21^+^ chondrocytes influences the extent of compensatory proliferation. By using a new *Col2a1-tTA* line (*Pit-tTA-p21* model), we misexpressed *p21* in fewer left limb chondrocytes (30-40% at E15.5, 15-35% at E17.5, 10-20% at P0, Supplemental Fig. 3A). Compensatory proliferation was not triggered (Supplemental Fig. 3B-C), suggesting it requires a minimum insult threshold. Lastly, we calculated the correlation coefficient between the % of p21^+^ chondrocytes and the extent of proliferation in GPs from left and right *ePit-Col-p21* (*in vivo* and *ex vivo*) and *Pit-tTA-p21* tibiae, at E17.5 (or E15.5+2days *ex vivo*). Segmental linear regression analysis revealed that the extent of EdU incorporation by p21^−^ chondrocytes did not correlate with the proportion of p21^+^ neighbours when this proportion was below 35%, but beyond this threshold, there was linear correlation between both parameters (Fig. 3D). These results suggest that compensatory proliferation is due to a signal produced in proportion to the number of arrested chondrocytes, that the signal needs to reach a certain threshold to be effective, and that it remains active until at least P0 despite the dilution of p21+ chondrocytes. Interestingly, we observed a temporal association between the occurrence of compensatory proliferation in the *ePit-Col-p21* model (i.e. at E17.5 and P0 but not E15.5) and statistically significant reduction of cell density in the left PZ as compared to the right (Fig. 3E). Notably, left and right PZ cell densities were not significantly different at any stage in *ePit-p21* mice (Fig. 3E, n=12). These findings raise the possibility that the signal triggering increased proliferation is related to the decreased cell density that follows chondrocyte arrest. In fact, we found that at E17.5 there was a threshold value of cell density below which EdU incorporation sharply increased in p21^−^ chondrocytes (Fig. 3F), likely explaining why a certain extent of insult is needed to trigger compensatory proliferation. Such a mechanism would also ensure compensatory proliferation does not lead to overgrowth once the threshold cell density is attained.

**Figure 3.**
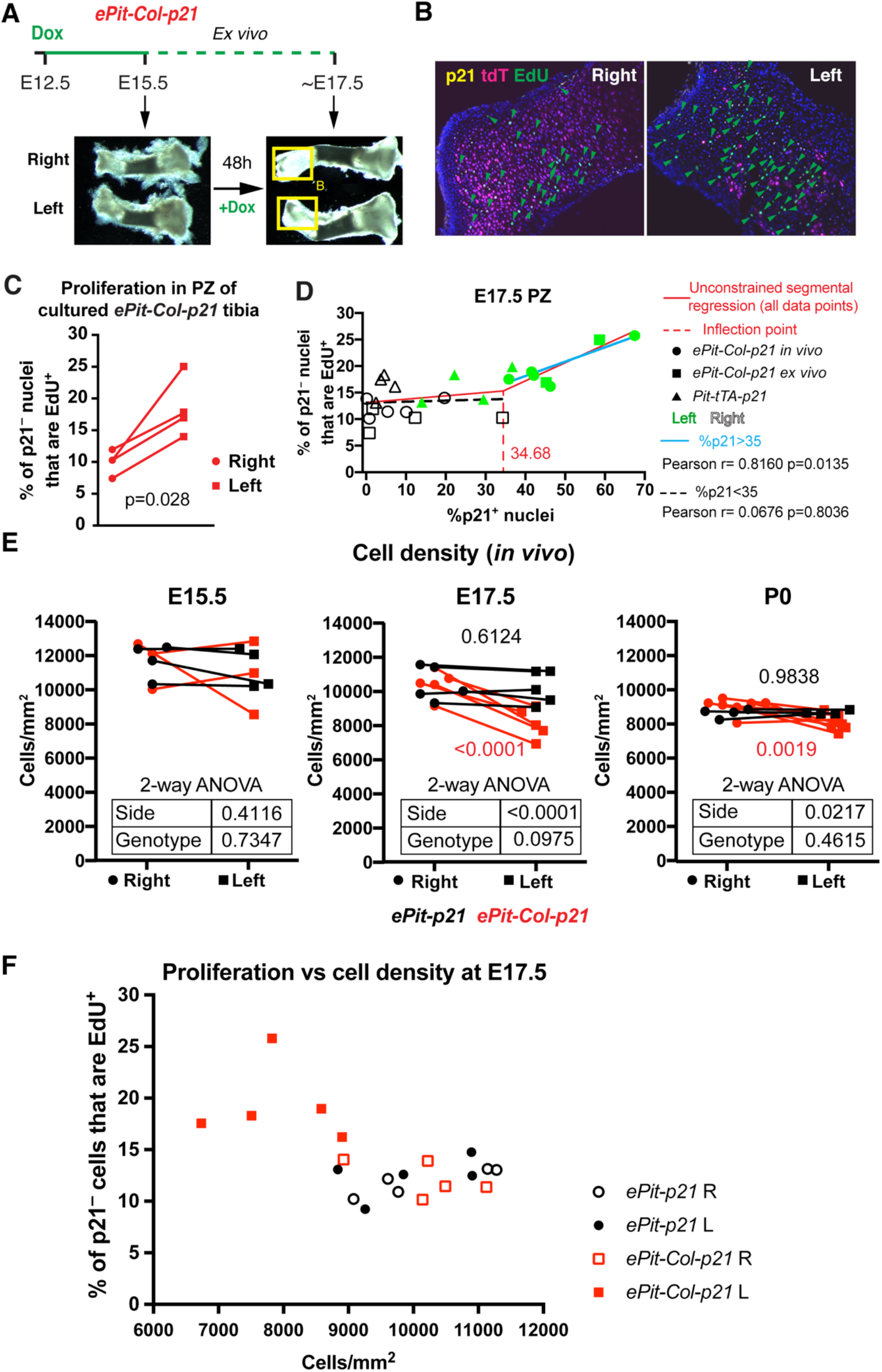
Compensatory proliferation takes place when cell density in the growth plate is lower than normal. (**A**) Summary of the *ex vivo* tibial culture experiment. The boxed regions correspond to the GPs shown in (B). (**B**) Immunohistochemistry for the indicated molecules (arrowheads= EdU^+^ chondrocytes). (**C**) EdU quantification on distal GP sections obtained from E15.5 *ePit-Col-p21* tibiae cultured for two days. p-value for two-tailed paired t-test comparing left and right proliferative ratios of p21^−^ chondrocytes is shown (n=4). The distal GP was quantified because the proximal one (bulkier) shows proliferation only in the periphery. (**D**) Correlation analysis between the extent of EdU incorporation in p21^−^ cells and the amount of p21^+^ nuclei in left and right PZ of *ePit-Col-p21* (n=5 *in vivo* and 4 *ex vivo*) and *Pit-tTA-p21* GPs (n=4) at E17.5. The inflection point revealed by unconstrained segmental regression was rounded up and used as a dividing threshold for the two correlation analyses (colour-coded). Pearson correlation coefficients and two-tailed p-values are shown. (**E**) Comparison of chondrocyte density in the proliferative zone (PZ) of left and right *ePit-p21* and *ePit-Col-p21* tibial GPs at E15.5 (n=4 and 3), E17.5 (n=5 and 5) and P0 (n=4 and 7) and analysed by 2-way ANOVA for Genotype and Side (p-values shown in the embedded tables). When p<0.05 for these variables, Sidak’s post-hoc tests are shown. (**F**) EdU incorporation in p21^−^ chondrocytes of left and right PZ from E17.5 *ePit-p21* and *ePit-Col-p21* embryos (n=5 each), plotted against cell density in the PZ. Note the sharp change in proliferation beyond 9,000 cells/mm^2^.

### Mosaic local proliferation blockade in chondrocytes of the left limb results in systemic growth reduction

Since the proliferative capacity of chondrocytes could have an intrinsic limit, we next tested whether systemic effects contribute to rescuing the induced growth defect. We indeed found that right bone length and body weight of E17.5 and P0, but not E15.5 *ePit-Col-p21* mice were ~10% lower than those of *ePit-p21* littermates, an effect that required Dox treatment and therefore *p21* expression (Fig. 4A-C, Supplemental Fig. 4A and not shown). Importantly, there was no leakiness of the intersectional misexpression strategy (Supplemental Fig. 4B) that could account for the systemic growth reduction, and misexpression of tdT in all chondrocytes does not cause a systemic growth reduction (Fig. 4B-C). Our results thus revealed a whole-body response to a local insult in mice, similar to what has been described in *Drosophila* larvae[2–5]. To characterize the cartilage response, we performed an RNA-seq experiment to identify differentially expressed genes (DEG) between left and right *ePit-Col-p21* GPs at E17.5 (Supplemental Fig. 5A-E). Indeed, overrepresentation analysis of the DEG (padj≤0.05) showed enrichment of several pathways related to stress and immune responses in the left cartilage (Supplemental Fig. 5F). In particular, we found several stress-related transcripts that shared a similar left-right pattern of expression within each embryo (Supplemental Fig. 5G) and verified their enrichment in the left cartilage by qRT-PCR (Fig. 4E) or *in situ* hybridisation (Fig. 4F). *Relaxin1*, the closest homologue to *dilp8*, the recently identified[3,22] alarm gene in fly, was not expressed at significant levels in either limb (Supplemental Fig. 5E), suggesting the mechanisms that link the local insult with a systemic response have diverged during evolution. Regarding the relationship between the extent of insult and the systemic response, *Pit-tTA-p21* mice did not trigger a systemic growth defect at E17.5 or P0 (Supplemental Fig. 3D-E, summary in Fig. 5A), suggesting that the systemic growth reduction is also only triggered when a certain insult threshold is surpassed in the targeted cartilage.

**Figure 4.**
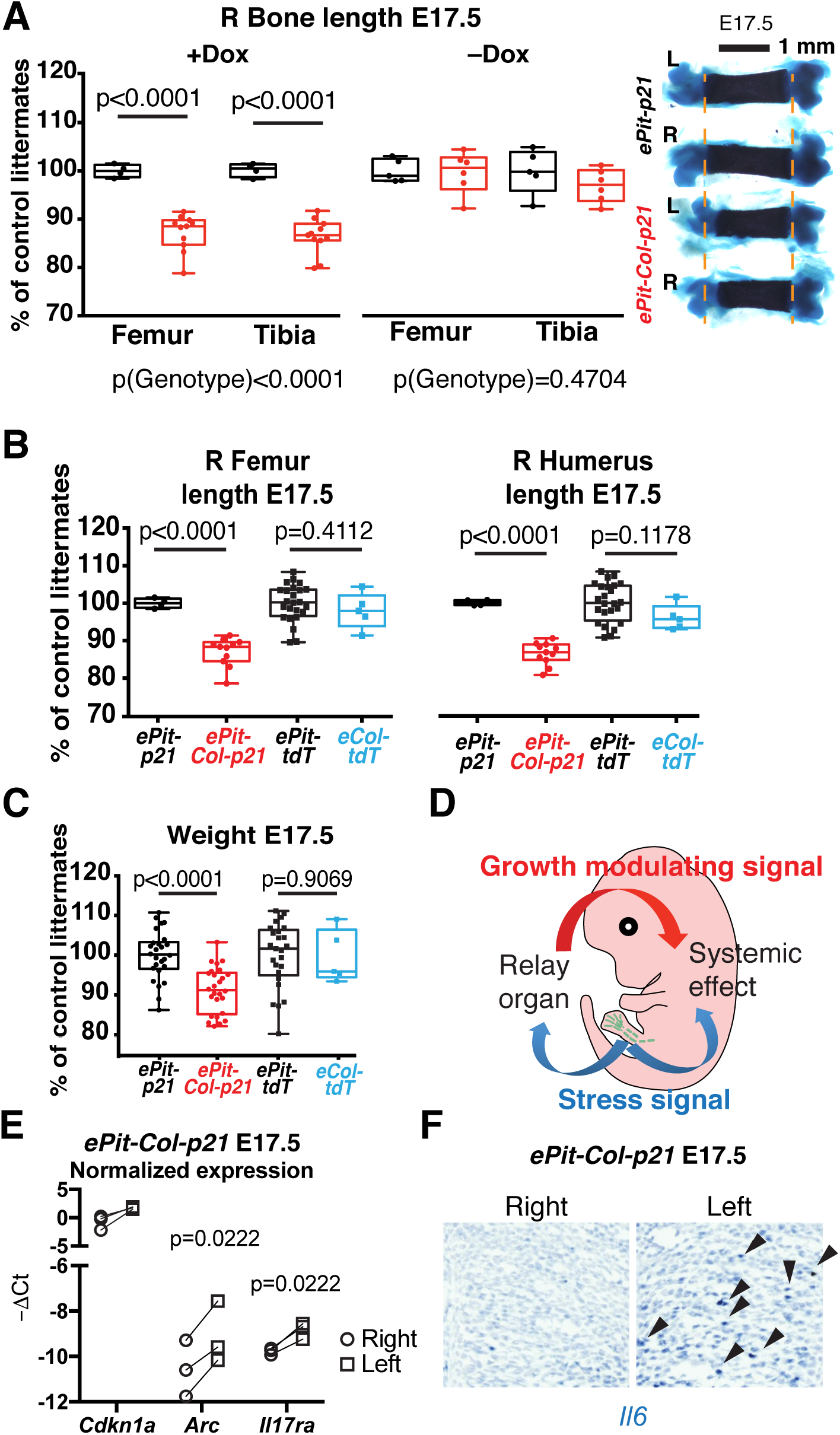
Mosaic local proliferation blockade in chondrocytes of the left limb results in systemic growth reduction. (**A**) Right femur and tibia length (normalised to the average *ePit-p21* littermate) from E17.5 embryos treated with Dox (n=4 *ePit-p21* and 11 *ePit-Col-p21*) or untreated (n=5 and 6). Comparison by 2-way ANOVA with Genotype and Bone identity as variables. p-values for Genotypes are shown below graphs, p-values for Sidak’s post-hoc test shown on graph. Femoral skeletal preparations are shown on the right (dashed lines flank the ossified region in control bones). (**B-C**) Box and whiskers plots for normalised bone length (B) and weight (C) of *ePit-Col-p21* and *Col2a1-rtTA; Igs7^TRE-tdT/+^* embryos (*eCol-tdT*, expressing tdT in all cartilage elements), compared by multiple unpaired t-tests. p-values corrected for multiple comparisons (Holm-Sidak method) are shown. For (B), n=22 *ePit-p21*, 26 *ePit-Col-p21*, 25 *ePit-tdT* and 5 *eCol-tdT*). For (C), n=4, 11, 24 and 5. (**D**) Model of the systemic growth response after local chondrocyte arrest triggers an alarm signal. (**E-F**) qRT-PCR (E) and *in situ* hybridisation (F) for the indicated transcripts in GPs from *ePit-Col-p21* embryos. (E) shows one of two independent experiments with 3 distinct biological replicates each (n total=6). The –ΔCt (relative to *Gapdh*) for each stress-related transcript was compared by a paired t-test (left vs. right). In (F), n=2 E15.5, 4 E16.5 and 6 E17.5 embryos (arrowheads denote *Il6* expression).

**Figure 5.**
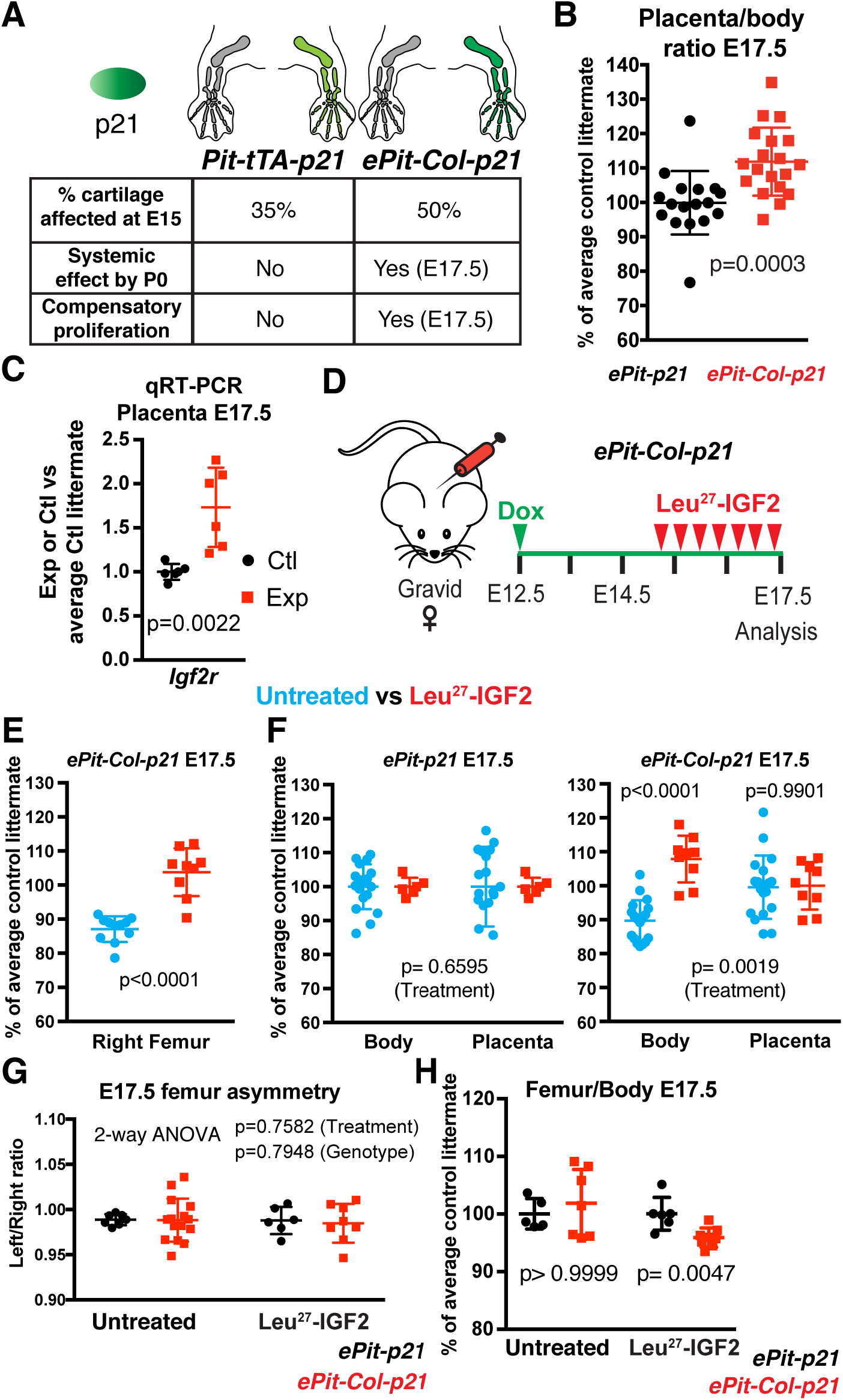
The systemic growth reduction of *ePit-Col-p21* embryos involves impaired placental function and is necessary to maintain limb/body proportions. (**A**) Summary of the characteristics and outcomes of the different injury models. Colour gradients reflect the extent of insult. (**B**) Weight ratio of the *ePit-Col-p21* placenta (n=19) with respect to the body, normalised to the average of *ePit-p21* littermates (n=17) at E17.5, and compared by two-tailed unpaired Mann-Whitney test. (**C**) qRT-PCR for *Igf2r* (with *Tbp* as reference gene) in the placenta of E17.5 *ePit-Col-p21* and *ePit-p21* embryos (n=6 each), normalised to the average value of control littermates. p-value for unpaired Mann-Whitney test is shown. (**D**) Pregnant females were treated with Leu^27^-IGF2 to improve placental efficiency. (**E-F**) Characterization of the systemic growth reduction at E17.5 in Leu^27^-IGF2-treated and untreated litters. For femur length, n=11 untreated and 9 treated *ePit-Col-p21* embryos. Unpaired two-tailed Mann-Whitney test was used. For body and placental weight, n=17 untreated and 6 treated *ePit-p21* embryos; 19 and 9 *ePit-Col-p21* embryos. 2-way ANOVA with Conceptus part and Treatment as variables was used. p-values for Treatment (bottom) and for Sidak’s post-hoc tests (top) are shown. (**G**) Left/right ratio of femur length for E17.5 *ePit-p21* and *ePit-Col-p21* embryos from Leu^27^-IGF2-treated (n=6 and 9) and untreated litters (n=4 and 11). p-values (2-way ANOVA) for Treatment and Genotype are shown. (**H**) Similar to (G), showing femur length/body weight ratio of E17.5 *ePit-p21* and *ePit-Col-p21* embryos, normalised to the average control littermate (n=5 and 7 untreated, 6 and 9 treated). For each treatment, comparisons by unpaired Mann-Whitney test are shown.

### The systemic growth reduction of ePit-Col-p21 embryos involves impaired placental function and is necessary to maintain limb/body proportions

We reasoned that the most likely organ to respond to a circulating alarm signal is the placenta, as in rodents it produces higher insulin-like growth factor (IGF) levels than any other organ[23] and is considered the main organ controlling foetal growth[24], whereas hepatic IGFs regulate systemic growth mainly after weaning[25]. Interestingly, placental weight was not diminished in *ePit-Col-p21* embryos as compared to *ePit-p21* controls (Supplemental Fig. 6A), such that the placenta/body weight ratio was increased (Fig. 5B). These results suggest that placental efficiency is reduced in response to the cartilage insult. Indeed, levels of *Igf2r* mRNA, which encodes a decoy receptor that decreases local IGF2 bioavailability[26], were increased in the placenta of *ePit-Col-p21* embryos compared to *ePit-p21* controls (Fig. 5C). To test if the systemic growth reduction in *ePit-Col-p21* embryos was due to impaired placental IGF signalling, we injected pregnant dams with an IGF2 analogue that can only bind IGF2R, and was thus expected to increase bioavailability of endogenous IGF2 and placental efficiency[27]. Confirming a role for placental function, *ePit-Col-p21* body weight and right femur length were significantly rescued in the treated litters, whereas placental weight remained unchanged (Fig. 5D-F and Supplemental Fig. 6B-D). Rescue of the systemic effect did not, however, result in left-right asymmetry in *ePit-Col-p21* embryos (Fig. 5G). However, the femur/body weight ratio of rescued *ePit-Col-p21* embryos was diminished compared to *ePit-p21* littermates or untreated litters (Fig. 5H). These results suggest that a decrease in growth of the unmanipulated limb contributes to the maintenance of left-right symmetry upon a unilateral insult, and that systemic growth reduction is therefore necessary to maintain limb/body proportions. Furthermore, the fact that left and right limbs are equally reduced in length suggests there is direct left-right crosstalk between the limbs, as previously proposed in studies on amphibian regeneration[28] and tibial fracture healing in young rats[29].

### A holistic view of the compensatory responses triggered by developmental insults

Collectively, our results reveal that the processes leading to coordination of growth within and between organs to achieve normal proportions upon developmental insults are conserved across metazoans. We propose that when an organ experiences developmental or environmental perturbations, an adaptive growth response that involves cell-nonautonomous local mechanisms interacting with systemic changes is initiated during the insult time frame to ensure that body proportions are maintained (Supplemental Fig. 7). The magnitude of the contributions of local and systemic mechanisms likely varies across phyla, however, as the extent of the systemic growth reduction observed in mice seems to be less extreme than in *Drosophila*. Finally, we speculate that the same ‘alarm’ signal triggers both the intrinsic and systemic mechanisms following injury, which would provide an evolutionary advantageous strategy to achieve robust coordination of organ growth. Further exploration of the mechanisms underlying these phenomena will open new exciting avenues for basic and translational research, and lead to a better understanding of human growth disorders.

## Methods

### Study Design

For each experiment, the minimum sample size was estimated using an online tool (http://powerandsamplesize.com/Calculators), based on the average SD observed in pilot experiments, to achieve an effect size of 3% (left/right bone length ratio), or 10% (rest of parameters), with a power of 0.8 and a 95% confidence interval. In Fig. 4B-C, two embryos (one from the *ePit-Col-p21* and one from the *eCol-tdT* populations) were abnormally small, possibly dead, and were excluded from the analysis. For comparison of qualitative expression, a minimum of two specimens per stage and five across several stages were used. The investigator measuring bone length was blinded to the treatment/genotype of the specimens. No blinding was done for other measurements. No randomization was used for animal processing.

### Statistics

When data were available for control and experimental, a normalised measurement (left/right ratio or % of average control mice) was calculated for both. Between different time points, the normalised measurements were compared by multiple unpaired t-test with Holm-Sidak correction for multiple comparisons. Within the same time point, comparisons were done by an unpaired Mann-Whitney test (one variable and two conditions), or by one-way ANOVA (one variable and ≥3 conditions) or by 2-way ANOVA (two variables and two or more conditions), following a matched (paired) design when possible. When left and right measurements were compared within experimental animals only, paired two-tailed t-test was used. For all ANOVA, alpha=0.05. All relevant parameters for the statistical tests can be found on Supplemental Table 3. When parametric tests were used, data normality was confirmed by Shapiro-Wilk test, and equality of variance by F-test. Prism7 software (Graphpad) was used for most analyses. Most graphs show individual values and mean±SD, unless otherwise indicated.

### Mice

To generate the *Igs7^TRE-LtSL-p21^* mouse line, the *NruI*-STOP-loxP-tdTomato-*SnaBI* fragment in the *Ai62*(*TITL-tdT*) *Flp-in* replacement vector[17] was replaced by a custom *NruI*-tdTomato-STOP-loxP-*MluI-HpaI-SnaBI* cassette, to generate an empty DRAGON vector. A PCR-amplified Kozak-Cdkn1a cassette was subsequently cloned into the *MluI* and *SnaBI* sites to generate the *DRAGON-p21* vector. This vector was then used for recombinase-mediated cassette exchange into Igs7-targeted G4 ES cells[17]. Two successfully targeted clones were injected into C2J blastocysts to generate chimeras, obtaining 27 chimeric males (out of 30 born) with 75-100% chimaerism. Two males from each clone were crossed to Black Swiss mice (Charles River) to assess germline transmission, and to establish the new mouse lines. To generate the *Col2a1-tTA* line, a *Kozak-tTA* fragment was PCR-amplified from plasmid *pEnt L1L3 tTA-3* (Addgene #27105) and cloned into a vector containing the regulatory region of mouse *Col2a1* obtained from plasmid *p3000i3020Col2a1* (ref. [30]). Backbone-free vector DNA was injected into FVB zygotes to generate transgenic lines. Four out of 11 founders transmitted the *Col2a1-tTA* allele. The progeny of one of those (founder #92) expressed the tTA faithfully in the highest percentage of chondrocytes, and was bred with *Pitx2-Cre* animals to generate breeders for the experiments. *Col2a1-tTA* mice were maintained on an outbred Swiss Webster background and genotyped using primers *Col2a1-F* (CCAGGGTTTCCTTGATGATG) and *tTA-R* (GCTACTTGATGCTCCTGATCCTCC) and a standard PCR program with 55°C annealing temperature. The Pitx2-Cre [19] (kind gift of Dr. H. Hamada), *Col2a1-rtTA[20]* (kind gift of Dr. K. Posey), *Ai9* (*R26^CAGGS-LSL-tdTomato^*)[31] and *Ai62* (*Igs7^TRE-LSL-tdTomoto^*)[17] mouse lines were maintained on an outbred Swiss Webster background and genotyped as previously described. *Igs7^TRE-LtSL-P21^* animals were genotyped like *Ai62* mice. *Pitx2-Cre/Cre; Col2a1-(r)tTA*/+ females and males homozygous for the conditional misexpression allele were crossed to generate experimental and control animals. Noon of the day of vaginal plug detection was considered E0.5. The equivalent of E19.5 is referred to as P0.

### Doxycycline treatment

Doxycycline hyclate (Sigma) was added to the drinking water at a final concentration of 1 mg/ml, with 1% sucrose to increase palatability.

### Leu^27^-IGF2 injections

Human Leu^27^-IGF2 (GroPep) was prepared at 500 ng/μl in sterile 0.01N HCl solution and kept at 4°C in between injections. From E15.25 to E17.25, the pregnant dam was subcutaneously injected every 8 hours, for a total dose of 1 μg/g of bodyweight per day.

### Skeletal preparations and measurements

Staining of cartilage and bone was performed as described[32]. For young mouse pups (≤P5), bone length was measured on digital microphotographs using the Line tool in Adobe Photoshop. Unless otherwise indicated, only the ossified region was measured. For adolescent and adult mice, the limbs were dissected out, skinned and incubated for a controlled time in 2% KOH at 37°C to remove the soft tissues. Individual bones were then measured using digital callipers (EZCal from iGaging). Tibiae were measured from the intercondylar eminence to the distal articular surface, while femora were measured from the trochanteric fossa to the intercondylar fossa.

### Sample processing for histology

Mouse embryos were euthanized by hypothermia in cold PBS. Mouse pups were euthanized by decapitation after hypothermia-induced analgesia. Knees (or isolated full tibiae and femora) were dissected out, skinned and fixed by immersion in 4% paraformaldehyde (PFA, Electron Microscopy Sciences) in PBS for 2 days at 4°C. After several washes with PBS, the tissue was then cryoprotected first by brief incubation with a solution of 15% sucrose and then 30% sucrose in PBS for at least 4 hours at 4°C, and then embedded in Cryomatrix (Thermo) using dry-ice-cold isopentane (Sigma). The knees were oriented sagittally and facing each other, with the tibiae on the bottom of the block (i.e. closest to the blade when sectioning). Serial 7-micron sections were collected with a Leica Cryostat on Superfrost slides, allowed to dry for at least 30 min and stored at -80°C until used. For all histological techniques, frozen slides were allowed to reach room temperature in a closed box, and Cryomatrix was washed away for 15 minutes with warm PBS (37°C).

### Immunohistochemistry and TUNEL

Sections were incubated in citrate buffer (10 mM citric acid, 0.05% Tween 20, pH 6.0) for 15 min at 90°C, allowed to cool down, washed with PBSTx (PBS containing 0.1% Triton X-100), blocked with 5% BSA in PBSTx 30 min at RT, and incubated with the primary antibody over night at 4°C (see list of antibodies below). After PBSTx washes, incubation with Alexa647- and/or Alexa555-conjugated secondary antibodies (Molecular Probes, 1/500 in PBSTx with DAPI) was performed for 1 h at RT. After PBSTx washes, the slides were mounted with Fluoro-Gel (Electron Microscopy Sciences). For TUNEL staining, endogenous biotin was blocked after antigen retrieval using the Avidin/Biotin blocking kit (Vector #SP-2001), and TdT enzyme and Biotin-16-dUTP (Sigma #3333566001 and #11093070910) were subsequently used following manufacturer instructions. Biotin-tagged DNA nicks were revealed with Alexa488- or Alexa647-conjugated streptavidin (Molecular Probes, 1/1000) during the incubation with the secondary antibody.

*Antibodies* (host species, vendor, catalogue#, dilution): tdTomato (rabbit polyclonal, Rockland #600-401-379, 1/500), p21 (rabbit polyclonal, Santa Cruz Biotechnology #sc-471, 1/300), p19^Arf^ (rat monoclonal, clone 12-A1-1, Novus Biologicals #NB200-169, 1/100).

### In situ hybridisation

The protocol described in[33] was followed. For embryos and young pups (P1-P5), samples were not decalcified. Except for *Col2a1, Col10a1* and *Ihh* (provided by Dr. Licia Selleri), the templates for most riboprobes were generated by PCR from embryonic cDNA, using primers containing the SP6 or T7 RNA polymerase promoters. Sequence of the primers is available upon request. After purification of the PCR product (Qiagen PCR purification kit), DIG-labelled probes were transcribed following manufacturer instructions (Roche), treated with DNAase for 30 min and purified by LiCl-mediated precipitation in alcoholic solvent. Probes were kept at - 80°C in 50% formamide (Fluka). For immunohistochemistry after *in situ* hybridisation, sections were incubated in citrate buffer (10 mM citric acid, 0.05% Tween 20, pH 6.0) for 15 min at 90°C, allowed to cool down, washed with PBSTx, and incubated with 1% H_2_O_2_ in PBSTx for 1 hour to block endogenous peroxidases. After BSA blocking and primary antibody incubation, endogenous biotin was blocked using Avidin/Biotin Blocking kit (Vector #SP-2001), and then the slides were incubated with a biotinylated secondary antibody. A brown precipitate was obtained using a peroxidase-coupled streptavidin-biotin complex (Vectastain Elite ABC Kit, Vector #PK-6100) and DAB substrate (Vector #SK-4100), following manufacturer instructions.

### Imaging

Bright-field and fluorescence images were taken on a Zeiss inverted microscope (Observer.Z1) using Axiovision software (Zeiss). Mosaic pictures were automatically reconstructed from individual 10x (brightfield) or 20x (fluorescence) tiles.

### EdU incorporation

5 mg/ml EdU in PBS was injected (50 μg/g body weight, s.c for pups, i.p. for adults and pregnant females) 1.5 h before euthanising the mice. EdU was detected using the Click-iT Alexa488 Imaging Kit (Invitrogen, C-10337), once the immunohistochemistry and/or TUNEL staining were finished on the same slides. The fraction of nuclei that were positive for EdU, p21 or tdTomato in the proliferative zone of the cartilage was determined using ImageJ.

### Cell size analysis

The proliferative zone was cropped from imaged sections of left and right *Pit-Col-p21* proximal tibial cartilage. tdTomato^+^ chondrocytes were segmented, measured and counted using Cell Profiler.

### RNA isolation and analysis

The distal left or right femoral and proximal tibial cartilage from E17.5 *Pit-Col-p21* embryos were dissected in cold PBS, the condyles and hypertrophic zones removed using a microknife, and the perichondrium removed by a combination of collagenase type II treatment (Worthington, 2mg/ml in DMEM, 2 min at room temperature) and mechanical dissection. Left and right cartilage fragments from each embryo (#1, 2 and 3) were kept in separated tubes and flash-frozen in liquid nitrogen. RNA was extracted using Trizol (Invitrogen) and a mechanical tissue disruptor.

### RNA-seq

High quality RNA was deep sequenced (≥50 million paired-end reads) by the New York Genome Center. Aligned reads were analysed using DESeq2 tool in R. A paired design was used, such that left and right comparison was performed for each specimen, which minimized the effect of sequencing batch and inter-specimen variability. Differentially expressed genes were obtained using a threshold of adjusted p-value ≤ 0.05. NMF library tools were used to generate heatmaps. Enrichment analysis was performed using DAVID[34] and WebGestalt[35]. *qRT-PCR*.

cDNA was synthesized from purified RNA using iScript reverse transcriptase (RT) as described by the manufacturer (Bio-Rad). Each target was amplified in triplicate, to obtain an average per sample, using SYBR Green (Applied Biosystems) on a StepOnePlus realtime PCR system (Applied Biosystems). Primer sequences are shown in Supplemental Table 4. Negative controls (no template) and no-RT cDNA controls were included for each primer/sample combination. Relative expression on each sample was calculated by the 2^-ΔCT^ method, with *Gapdh* (for cartilage) or *Tbp* (for placenta) as a reference.

## Study approval

All animal studies were performed under an approved Institutional Animal Care and Use Committee mouse protocol according to MSKCC institutional guidelines.

## Data availability

The datasets generated during and/or analysed during the current study are tabulated in the Supplemental Information and archived at the following databases: GSE97232.

## Acknowledgements

We thank the Joyner lab for scientific discussions, D. Stephen for technical support, N.S. Bayin, J.M. González-Rosa and A. Wojcinski for comments on the manuscript, the Mouse Genetics core for generating chimeras and new transgenic lines, B. de Crombrugghe for the *p3000i3020Col2a1* vector and H. Hamada and K. Posey for the *Pitx2-Cre* and *Col2a1-rtTA* mouse lines, respectively. This work was supported by grant R21HD083860 (NIH-NICHD) to A.L.J., HFSP and Charles Revson postdoctoral fellowships to A.R.D and an NCI Cancer Center Support Grant (P30 CA008748) to MSKCC.

## Author contributions

A.R.D. conceived the approach, proposed the hypotheses and performed most experiments. L.M. and H.Z. generated and provided the Igs7-targeted ES cells. S.B. helped with the characterization of the systemic growth reduction. A.R.D. and A.L.J. designed the study, interpreted results and wrote the manuscript. A.L.J. supervised the study.

## Author information

The authors declare that no competing financial interests exist.

## Supplemental Information

**Supplemental Table 1 (separate file)**

Normalised counts for deep-sequenced transcripts from left (L) and right (R) proliferative cartilage from three different E17.5 *ePit-Col-p21* embryos. The original numbering (#386-388) was changed to #1-3.

**Supplemental Table 2 (separate file)**

List of differentially expressed genes between left and right *ePit-Col-p21* cartilage at E17.5. The DESeq2 tool (padj ≤ 0.05) was used to obtain the list.

**Supplemental Table 3 (separate file)**

Parameters of the statistical tests used in this study.

**Supplemental Table 4.**
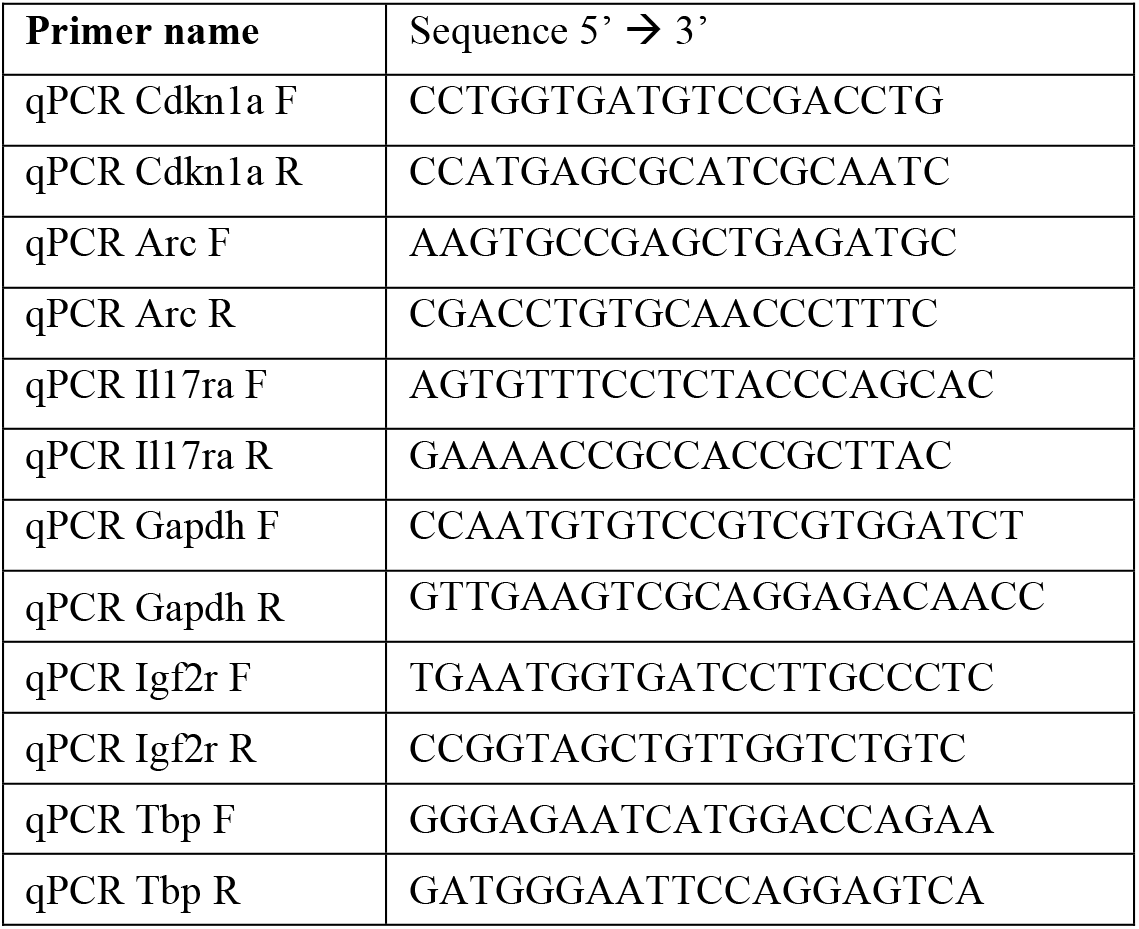
Sequence of the oligonucleotides used for qRT-PCR

**Supplemental Figure 1.**
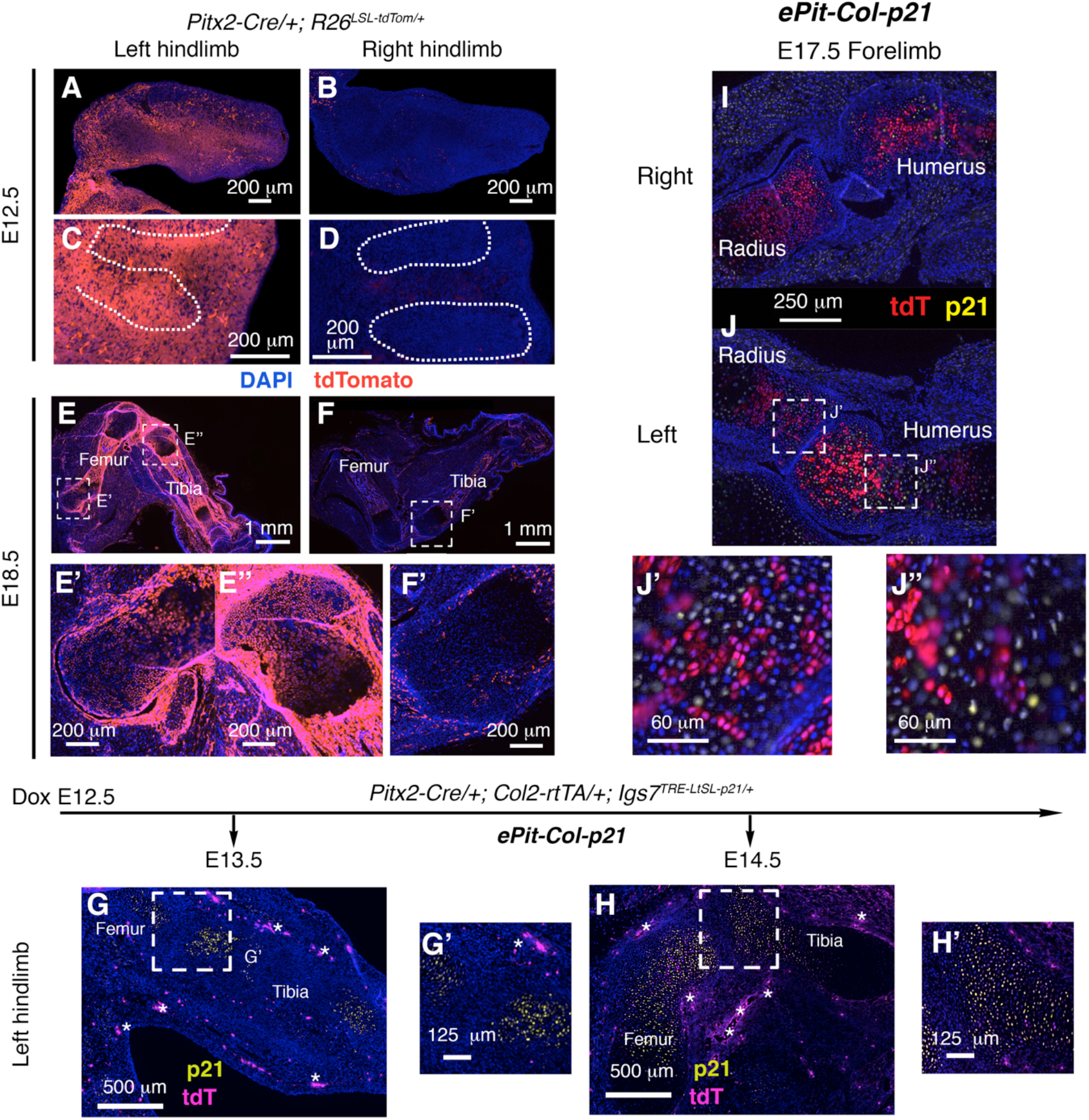
Characterization of the left limb-specific intersectional approach to induce transient growth defects. (**A-F’**) *Pitx2-Cre* females were crossed with Ai9 males to characterize the specificity of Cre-mediated labelling. 7-μm sections from left and right hindlimbs are shown at two different stages: E12.5 (A-D) and E18.5 (E-F’) n = 4 for each stage. Boxed regions in (E) and (F) are shown in (E’), (E’’) and (F’). Most of the red signal on right limbs corresponds to autofluorescent blood cells. (**G-H’**) Dynamics of tdTomato and CDKN1A (p21) activation in *ePit-Col-p21* embryos, one (G, G’, n=2) and two days (H, H’, n=3) after Dox administration to the pregnant female. Boxed regions in (G) and (H) are shown in (G’), and (H’). Note that activation of the transgene starts to be detectable one day post Dox administration, but it is not complete until two days post-Dox. Asterisks = autofluorescent cells. Of note, the *Pitx2-Cre* allele is consistently left-predominant only when inherited from the female (not shown). (**I-J”**) Same as above, but E17.5 forelimb sections are shown.

**Supplemental Figure 2.**
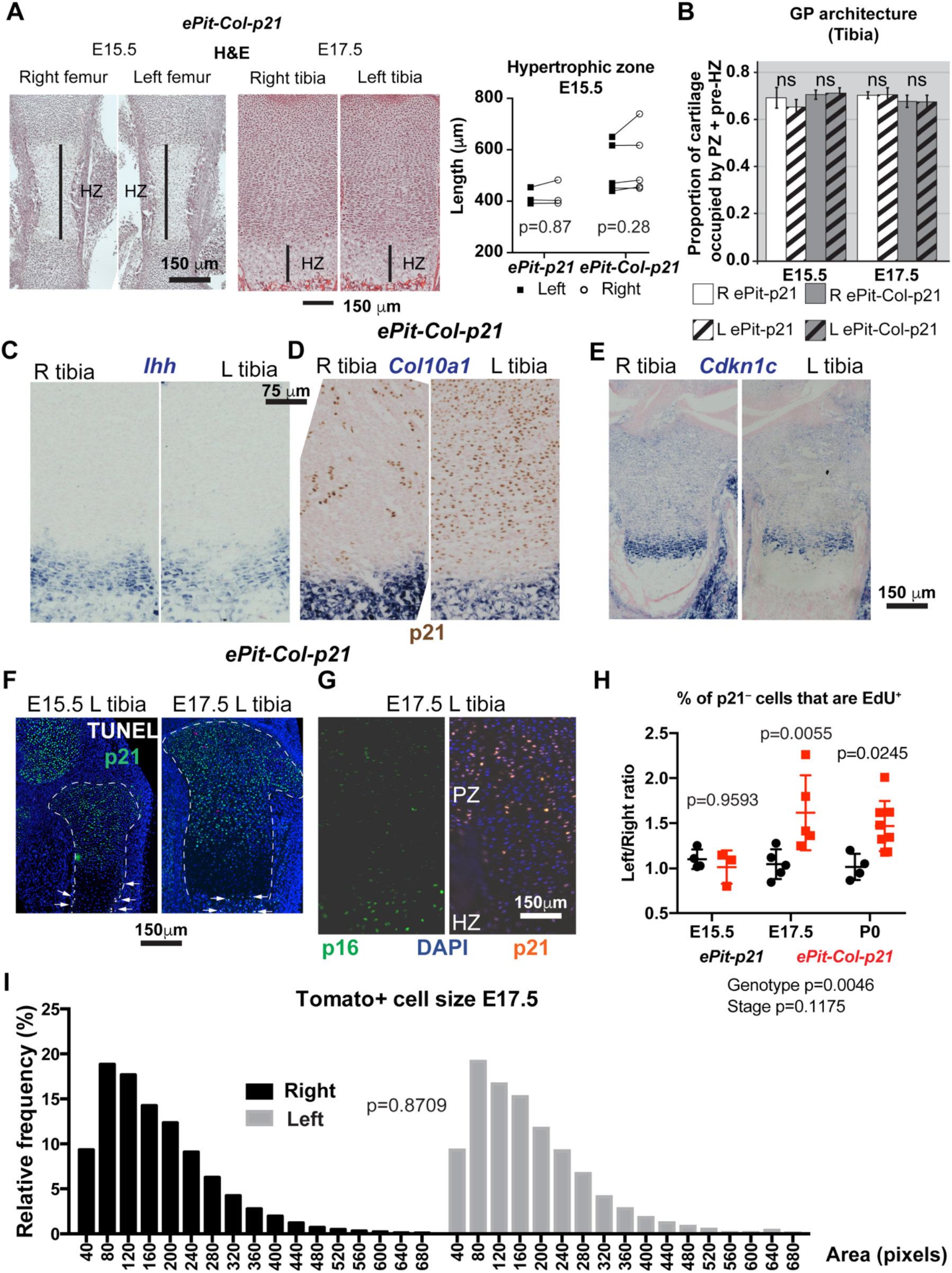
Histological, molecular and cellular characterization of the effects of *p21* misexpression. (**A**) Hematoxylin and eosin (H&E) staining of E15.5 femora and E17.5 proximal tibiae from *ePit-Col-p21* embryos and comparison of the length of the left and right hypertrophic zone (HZ) of the femora from *ePit-Col-p21* and *ePit-p21* embryos at E15.5 (2-way ANOVA with Genotype and Side as variables was used, and p-values are shown). (**B**) The stratification of the proximal tibial cartilage, expressed as the proportion of the total cartilage length represented by the sum of the proliferative zone (PZ) and pre-hypertrophic zones (pre-HZ), is not significantly different between left (L) and right (R) experimental and control embryos at E15.5 or E17.5 (n= 2 embryos for each genotype and stage). 2-way ANOVA with Genotype and Side as variables was used, and p-values for each stage are shown. (**C-E**) The expression of chondrocyte maturation markers *Cdkn1c, Col10a1* and *Ihh* is not ectopically triggered by p21 misexpression. (**F-G**) Misexpression of p21 does not lead to ectopic cell death in the experimental cartilage at E15.5 or E17.5 [(F), arrows, n=5) or cell senescence at E17.5 [(G), monitored by p16 expression, n=2]. (**H**) Left/right ratios of EdU+ incorporation in the PZ of *ePit-p21* and *ePit-Col-p21* embryos at E15.5 (n=4 and 3), E17.5 (n=5 and 5) and P0 (n=4 and 8). Comparison by 2-way ANOVA for Genotype and Stage (p-values below graphs). p-values for Sidak’s posthoc test are shown on the graphs. (**I**) Cell size of WT (tdT+) chondrocytes was characterized for *ePit-Col-p21* embryos at E17.5 (n= 10). No significant difference between left and right distribution was found (p-value for two-tailed unpaired Mann-Whitney test for ranks is shown).

**Supplemental Figure 3.**
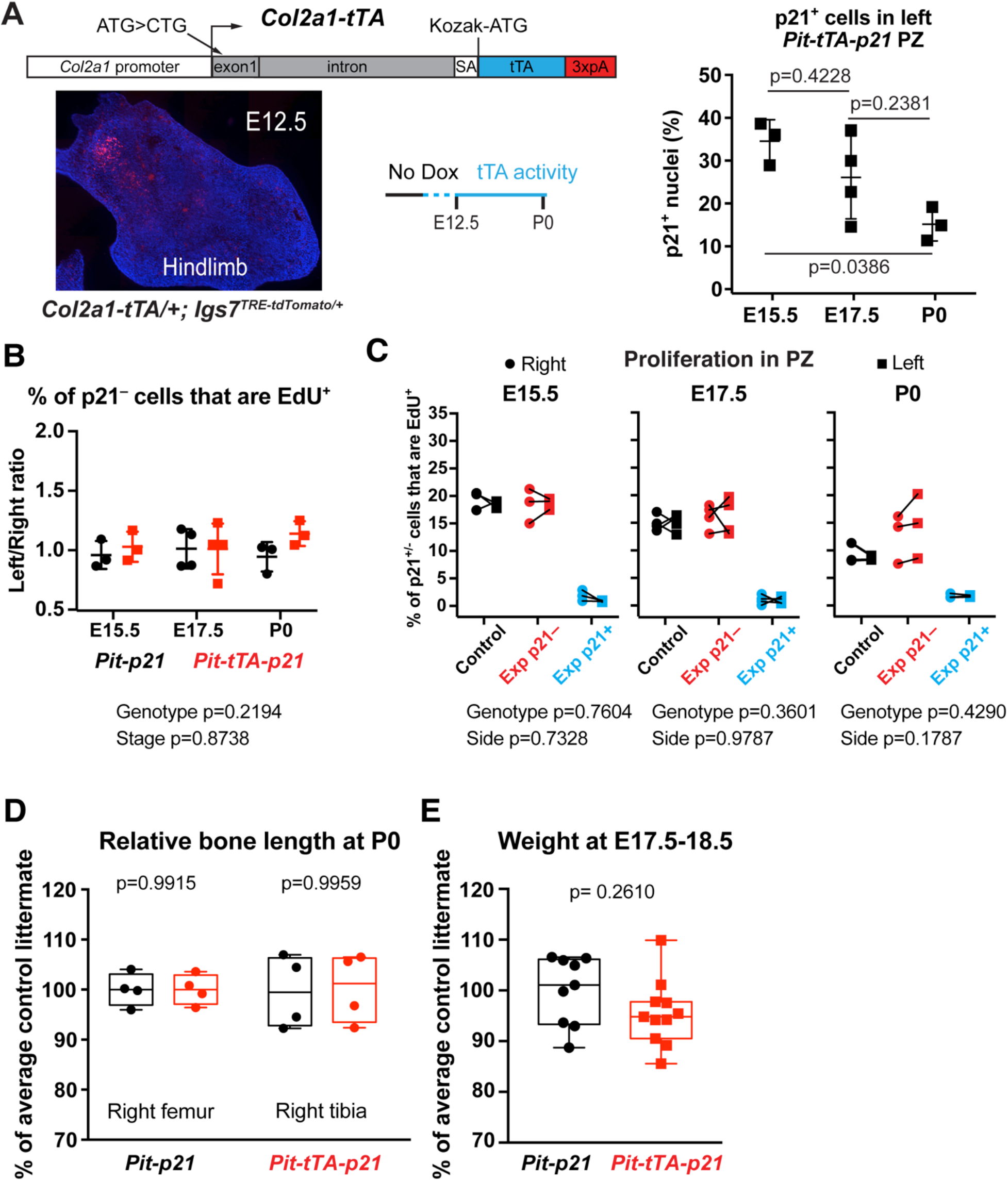
Compensatory proliferation and systemic growth reduction are not detected by birth when *p21* is expressed in less than 35% of chondrocytes. (**A**) Top: schematic of the new *Col2a1-tTA* allele. See ref. [30] for details on the *Col2a1* regulatory region used. In the absence of Dox, the tTA is activated around E12.5 (detected by a germline-recombined reporter *Ai62* allele)[17]. Right graph: % of p21+ chondrocytes in the PZ of left and right proximal tibial GP of *Pit-tTA-p21* embryos unexposed to Dox, at E15.5, E17.5 and P0 (n=3, 4 and 3). Comparison by one-way ANOVA (p=0.0368), followed by Tukey’s posthoc tests (shown). (**B**) Left/Right ratio of EdU incorporation in PZ chondrocytes of *Pit-tTA* and *Pit-tTA-p21* mice at E15.5 (n=3 each), E17.5 (n=4 each) and P0 (n=3 and 3). Comparison by 2-way ANOVA for Genotype and Stage (p-values below graphs). (**C**) % of p21+ or p21^−^ chondrocytes that have EdU+ nuclei in the proliferative zone (PZ) in the left and right tibias of E17.5 *ePit-p21* (Control) and *ePit-Col-p21* (Exp) embryos. p21^−^ cells from Control and Exp mice were compared by 2-way ANOVA with Side and Genotype as variables (p-values below graphs). n as in (B). (**D**) Length of P0 *Pit-p21* (n=4) and *Pit-tTA-p21* (n=4) right bones, normalised to the average value of control littermates. Comparisons were done by 2-way ANOVA with Genotype and Bone identity as variables, p(Genotype)= 0.9800, p-values for Sidak’s posthoc test are shown. (**E**) Weight of pooled E17.5 and E18.5 *Pit-p21* (n=9) and *Pit-tTA-p21* (n=11) mice, normalised to the average value of control littermates, and compared by unpaired two-tailed Mann-Whitney test.

**Supplemental Figure 4.**
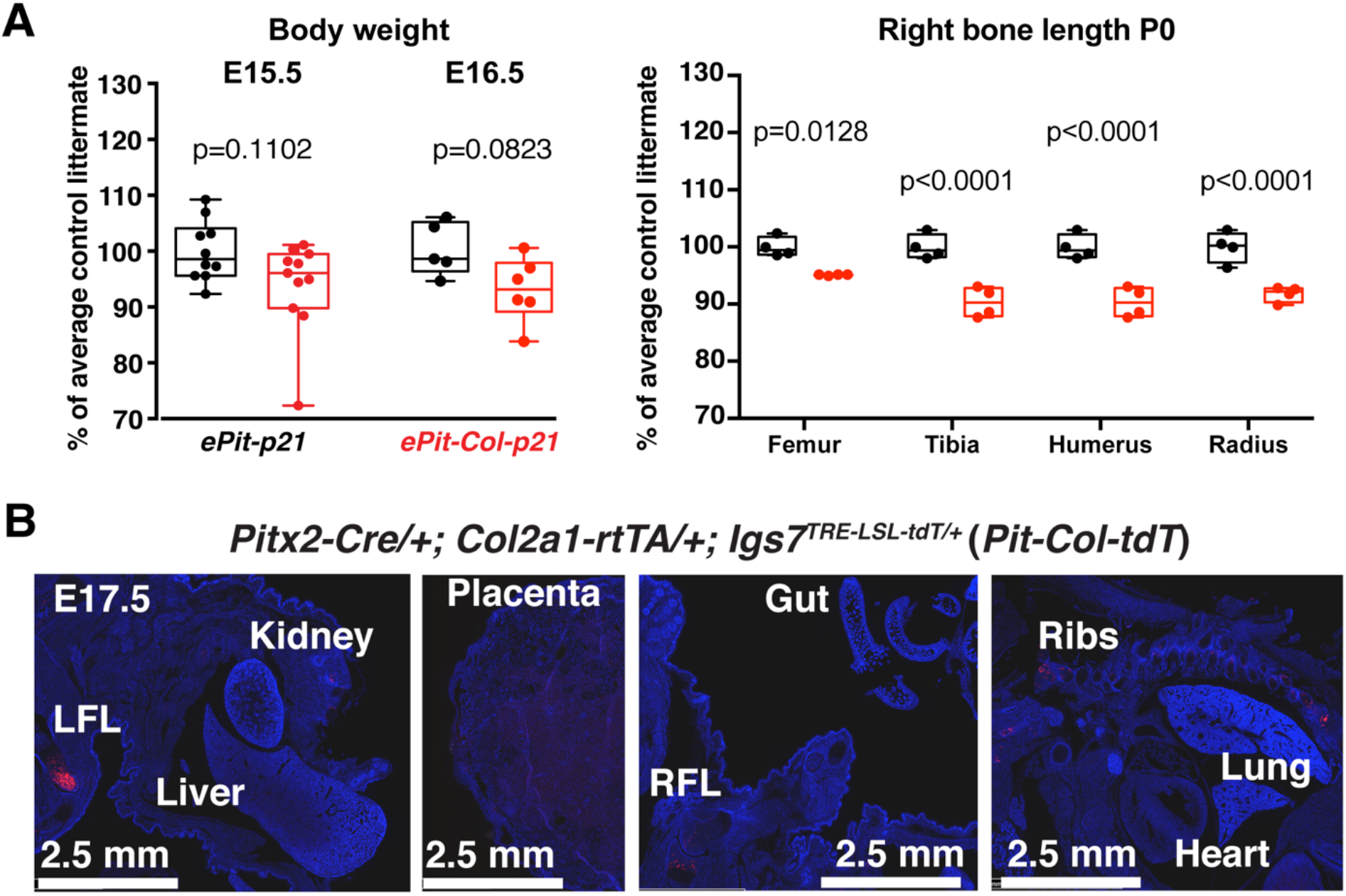
The systemic growth reduction triggered by transient and local p21 misexpression is progressive and not due to leakiness in other organs. (**A**) Left panel: Weight of E15.5 and E16.5 *ePit-p21* (n=10 and 5) and *ePit-Col-p21* (n=11 and 6) embryos, normalised to the average control littermate, and compared by two-tailed unpaired Mann-Whitney test. Right panel: comparison of right bone length at P0. n=4 *ePit-p21* and 4 *ePit-Col-p21* pups. Comparison by 2-way ANOVA with Bone and Genotype as variables. For Genotype, p=0.0004. p-values for Sidak’s posthoc test are shown. (B) Analysis of tdT expression in E17.5 *Pitx2-Cre/+; Col2a1-rtTA/+; Igs7^TRE-LSL-tdT/+^* embryos (*Pit-Col-tdT* model, Dox at E12.5) does not reveal spurious activation outside the left cartilage templates (n=2). LFL, RFL= left, right forelimb. The embryos were bisected sagittally to facilitate sectioning.

**Supplemental Figure 5.**
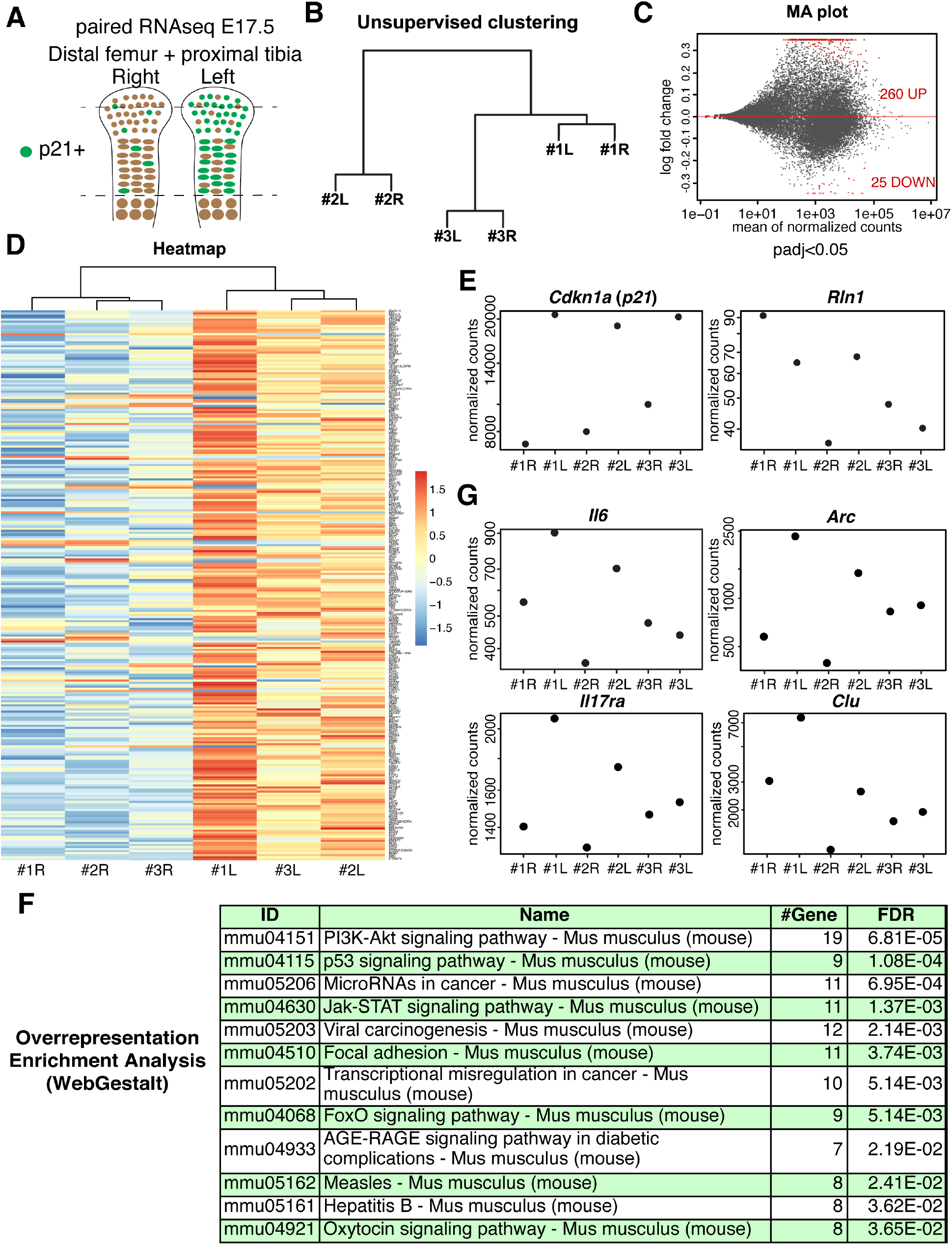
Transcriptomic comparison of left and right *ePit-Col-p21* cartilage. (**A**) Schematic of the experimental approach. After dissection and perichondrium (blue layer) removal, left and right cartilage elements were deprived of condyles and hypertrophic zone, and flash frozen. Left and right samples from each embryo were kept separated and RNA was extracted for deep sequencing. (**B**) Unsupervised hierarchical clustering of 6 samples (left and right cartilage from 3 embryos). Note that each sample is closest to its contralateral one. (**C-D**) MA plot (C) and clustered heatmap (D) of the 285 differentially expressed genes [red dots in (C)] obtained by a paired DESeq2 design with adjusted p-value ≤ 0.05. (**E**) Normalised counts for *Cdkn1a* (*p21*) and *Rln1* (*Relaxin1*, the closest vertebrate homologue to *dilp8*) are shown for each sample. Note that *Rln1* is virtually absent from control and experimental cartilage. See also Supplemental Tables 1 and 2. (F) Overrepresented pathways obtained from the 285 differentially expressed genes (FDR<0.05). Note the presence of immune response pathways. (**G**) Normalised counts for the transcripts following a similar left-right pattern as *Cdkn1a*. The four examples shown are involved in cellular stress response[38–41].

**Supplemental Figure 6.**
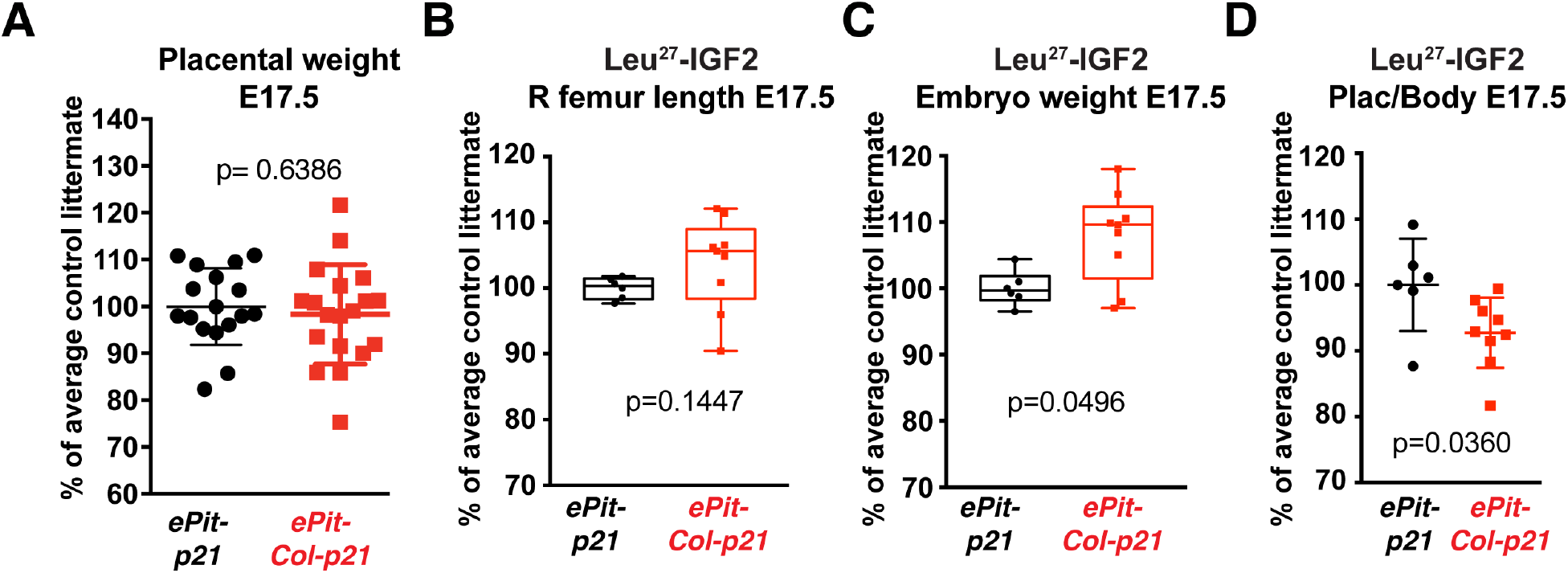
Reduced placental function underlies the systemic growth reduction in *ePit-Col-p21* embryos. (**A**) Placental weight for E17.5 *ePit-p21* (n=17) and *ePit-Col-p21* embryos (n=18), normalised to the average of *ePit-p21* littermates, and compared by two-tailed unpaired Mann-Whitney test. (**B-D**) Comparison of the indicated body measurements at E17.5 between *ePit-p21* (n=6) and *ePit-Col-p21* embryos (n=9) from Leu^27^-IGF2-treated litters. Unpaired two-tailed Mann-Whitney test was used in all cases.

**Supplemental Figure 7.**
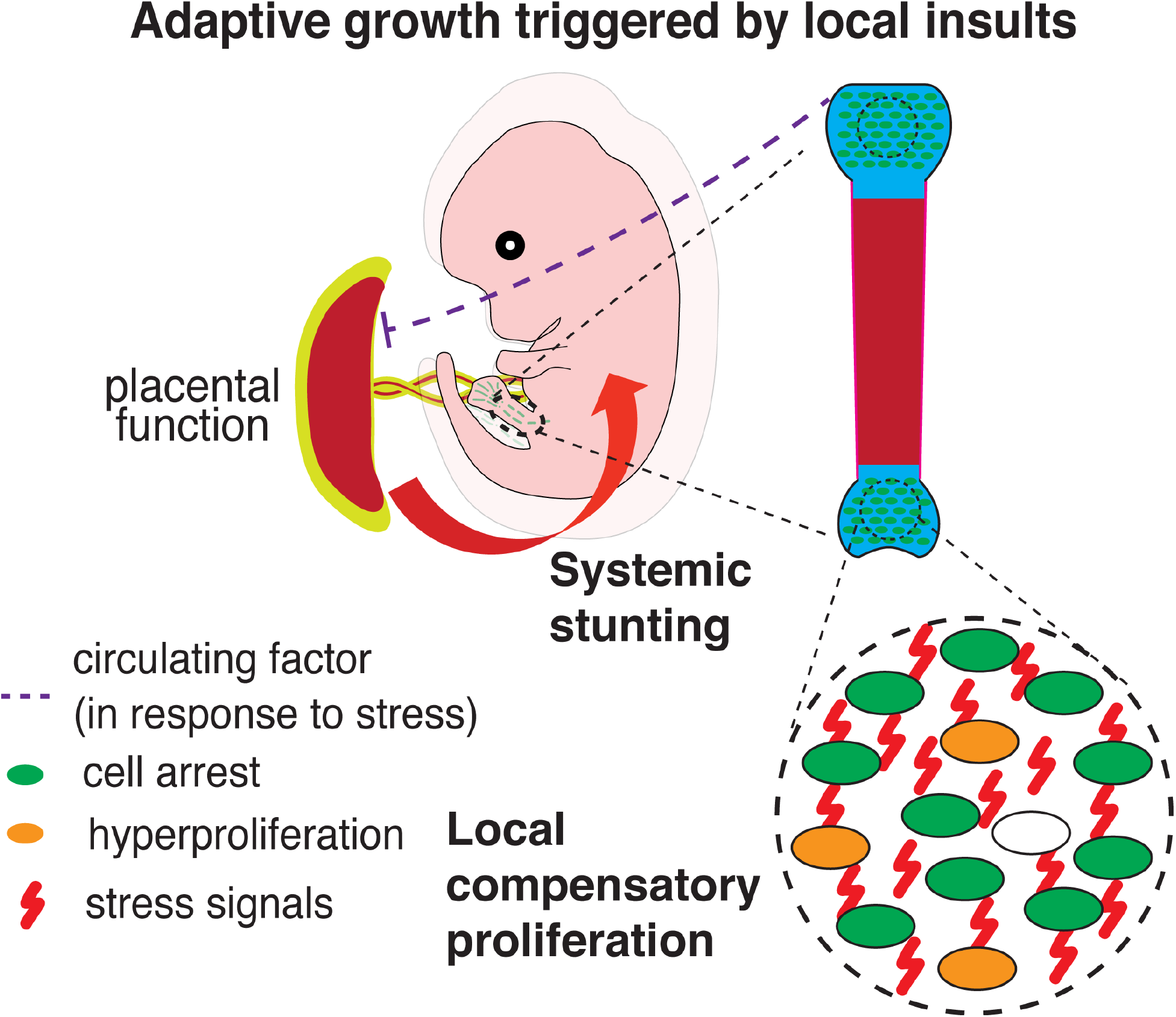
Model for adaptive growth after unilateral mosaic growth inhibition in long bone chondrocytes. Both local (compensatory proliferation) and systemic responses (reduced placental function) are triggered following expression of p21 in more than 35% of the growth plate chondrocytes in the left hindlimb.

